# Improve Protein Solubility and Activity based on Machine Learning Models

**DOI:** 10.1101/817890

**Authors:** Xi Han, Wenbo Ning, Xiaoqiang Ma, Xiaonan Wang, Kang Zhou

## Abstract

Improving catalytic ability of protein biocatalysts leads to reduction in the production cost of biocatalytic manufacturing process, but the search space of possible proteins/mutants is too large to explore exhaustively through experiments. To some extent, highly soluble recombinant proteins tend to exhibit high activity. Here, we demonstrate that an optimization methodology based on machine learning prediction model can effectively predict which peptide tags can improve protein solubility quantitatively. Based on the protein sequence information, a support vector machine model we recently developed was used to evaluate protein solubility after randomly mutated tags were added to a target protein. The optimization algorithm guided the tags to evolve towards variants that can result in higher solubility. Moreover, the optimization results were validated successfully by adding the tags designed by our optimization algorithm to a model protein, expressing it *in vivo* and experimentally quantifying its solubility and activity. For example, solubility of a tyrosine ammonium lyase was more than doubled by adding two tags to its N- and C-terminus. Its protein activity was also increased nearly 3.5 fold by adding the tags. Additional experiments also supported that the designed tags were effective for improving activity of multiple proteins and are better than previously reported tags. The presented optimization methodology thus provides a valuable tool for understanding the correlation between amino acid sequence and protein solubility and for engineering protein biocatalysts.

## Introduction

The exploration of expressing recombinant proteins started in 1976, when human peptide hormone Somatostatin was produced in *Escherichia coli*^1^. As the most commonly used expression host, *E. coli* was investigated intensively to improve the expression and activity of recombinant proteins^2, 3, 4^. Various experimental strategies, such as using protein fusion partners, co-expressing chaperones, choosing suitable promoters, optimizing codon usage, changing culture conditions, or using directed evolution^5, 6, 7, 8, 9, 10^, were used to improve protein expression. For example, the expression of human recombinant enzyme N-acetylgalactosamine-6-sulfatase (rhGALNS) in *E. coli* was undesirable due to protein aggregation. Several methods including the use of physiologically-regulated promoters, overexpression of native chaperones and applying osmotic shock were investigated to improve the production and activity of rhGALNS^10^. Protein activity, a phenotype representing the catalytic ability of a protein if it is an enzyme, is partly determined by its genotype (sequence of its coding gene). Directed evolution can effectively improve protein activity through changing the associated genotype, but this approach is resources-intensive. In the process of improving protein activity via directed evolution, mutagenesis is performed to change gene sequence and the mutated genes are inserted into plasmid used for transformation of a microbe, usually *E. coli*. Additional techniques are employed further to screen a large number of transformed cells for those that have higher protein activity. Since most of the protein directed evolution studies were only interested in the mutants with the highest activity, they did not reveal the genotype of most proteins that had lower activity. This fact has caused the challenge that almost no suitable database of protein activity is available for training computational models that can predict protein activity from protein sequence. Such models would greatly assist protein engineering by evaluating protein sequences *in silico*. A suitable dataset for training the model should contain both protein activity data and the associated sequence data, and should be large enough (>1,000 entries).

Protein activity data cannot be easily pooled together for model training if they are related to enzymes that catalyze different chemistries, which is another reason why it is difficult to generate the aforementioned datasets. The data of protein solubility from most types of proteins, however, can be compiled into one dataset, because protein solubility is a basic protein property. In this study and the relevant literature, protein solubility is defined as the percentage of a protein’s soluble fraction^11^. It is a metric that is often used to assess the folding quality of a protein, under the assumption that incorrectly folded proteins form aggregates and are insoluble. Protein activity is thus correlated with protein solubility to some extent, because protein solubility may indicate the quality of protein folding which influences protein 3D structure and activity, i.e. proteins with higher solubility likely exhibit higher activity^12^. Improving the solubility of some recombinant proteins can enhance their production effectively^13^. Thus, protein solubility may be used as a proxy for protein activity to develop predictive models that use protein sequence as input. With such a model, it would be possible to optimize the protein sequence of a protein *in silico* for improving its solubility and activity. For example, a Monte Carlo optimization method can be used as the procedures demonstrated in Figure 1: (1) a random change is introduced to the protein sequence, (2) the new protein sequence is evaluated by the model, and (3) if the predicted solubility is lower than that of the parental sequence, the change would be rejected, otherwise it would be accepted and used to initiate the subsequent iteration. This *in silico* optimization process may identify promising protein sequences to improve the success rate of the time-consuming and labor-intensive experiments. If the protein activity heavily depends on its solubility, the experiment would identify new protein that has higher solubility and activity.

**Figure 1.**
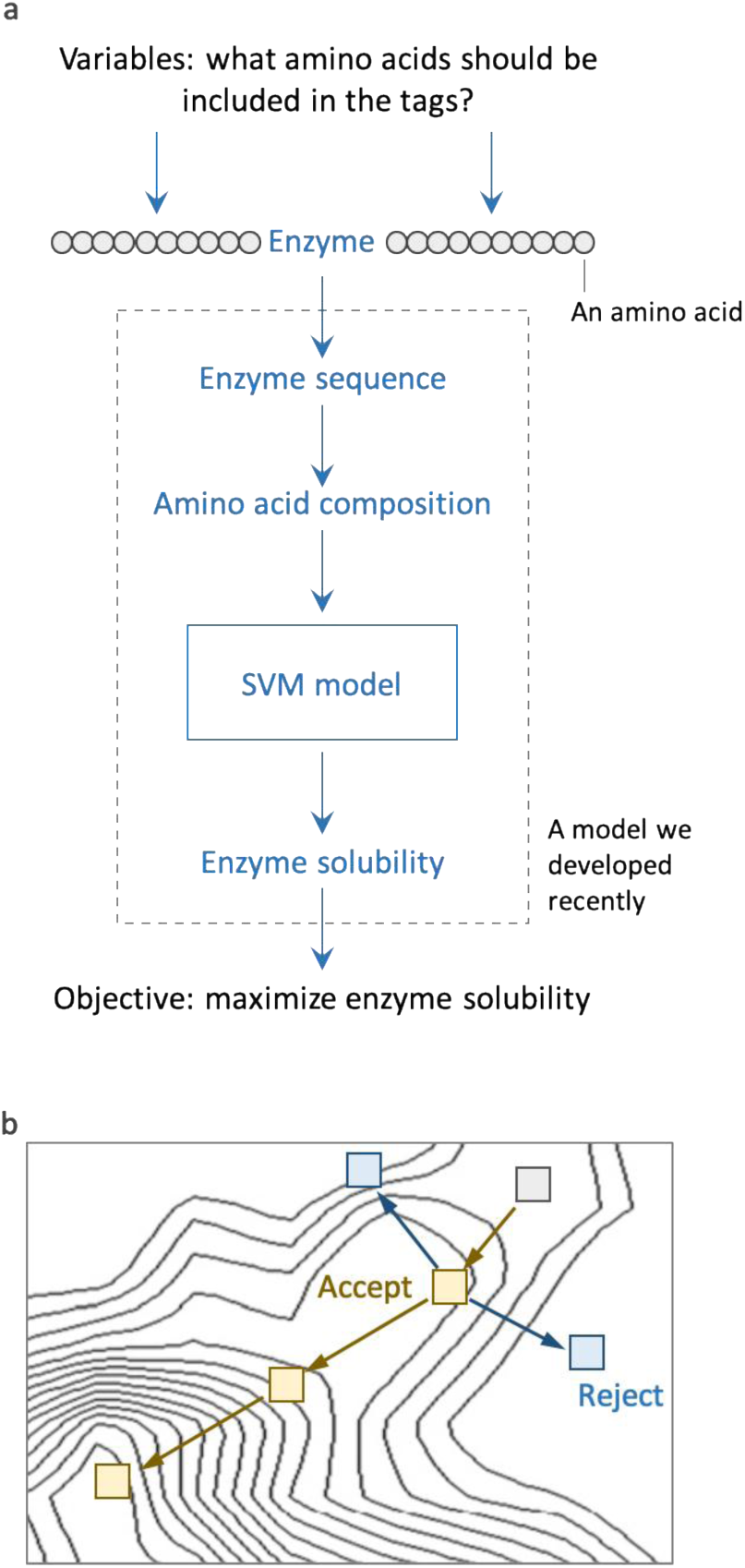
Machine learning model assisted optimization of protein solubility. (**a)** Illustration of the objective function when we aimed to improve protein solubility by adding short peptide tags. SVM: support vector machine. A SVM regression model we recently developed was used in this study^26^. (**b)** Illustration of the optimization algorithm. Genetic algorithm was used in this study.

Machine learning has gained increasing attention recently in various fields, such as internet commerce, autonomous vehicles, and image recognition^14, 15, 16, 17, 18, 19, 20, 21, 22^. Until now, a large number of machine learning methods have been explored to predict protein solubility from amino acid sequence^6, 11, 23, 24, 25^. Among the previous studies, we developed regression models that can predict protein solubility in the continuous values^26^. Classification models which only label a protein as soluble or insoluble were developed in other studies but cannot be used in the *in silico* optimization, because it would mistakenly reject most changes that can result in a small but important increase in the protein solubility. So far, very few studies performed experimental validation of their solubility-prediction models and no study used such models to improve protein properties through the *in silico* optimization of protein sequence.

In our present study, based on a regression model that can predict protein solubility from protein sequence^26^, we developed optimization algorithms to increase predicted solubility under constraints that have been set after considering experimental feasibility and impact on protein function. The performance of the optimization process for improving protein solubility was validated successfully by experimentally measuring solubility. We found that adding short peptide rich in negatively charged amino acids was effective in improving solubility of many proteins. More importantly, we also verified that activity of some proteins was indeed substantially improved when their solubility was increased. Our study provides a generally effective approach to enhance protein solubility and activity.

## Results

### Design the optimization methodology

In order to improve protein solubility by *in silico* mutagenesis, we need to solve several questions regarding how to change the protein sequence. One can change a protein sequence by adding amino acids to the sequence (addition), replacing amino acids in the sequence (mutation) and/or removing amino acids from the sequence (deletion). The protein functions may be frequently abolished by mutation and deletion as the original protein structure and active sites may be changed. To avoid such detrimental change to the original function of the protein, addition was used in our study to change protein sequence for improving protein solubility. The subsequent decision to make is how many amino acids should be added. Adding too many amino acids would make experimental validation to be more expensive and may also negatively affect the protein function. Adding too few amino acids may not be able to improve protein solubility substantially. We decided to evaluate adding 20 or 30 amino acids because adding more than 30 amino acids to a protein by using synthetic oligonucleotides was experimentally difficult.

To optimize the sequence of the amino acids to be added, we designed an algorithm based on the support-vector machine (SVM) prediction model we previously developed^26^. The independent variable in the optimization function is the amino acid composition of the short peptide to be added, expressed as number of each amino acid in a vector (Figure 1). The SVM model we developed only accepted amino acid composition of a protein as input, so we did not consider the full sequence information during the optimization. Then the amino acid composition of a model protein with the added amino acids was calculated and used as input for the SVM model. We used the genetic algorithm (GA) which is a widely used algorithm for solving constrained optimization problems. The objective function of GA outputs the predicted protein solubility by using the SVM model in the format of continuous values between 0-1. The sum of the number of amino acids added was set as 20 or 30 and the searching range for the number of each amino acid added was from 0 to 20 or 30.

### Optimize protein sequence *in silico* for improving protein solubility

After designing this optimization algorithm, ten proteins with low solubility (0.1) in the eSol database (we had used the same database to train our machine learning model) were selected as model proteins to test the algorithm (information of these proteins is provided in Supplementary Table S2). The predicted solubility of all the ten proteins was improved after adding 30 amino acids as peptide tags (Supplementary Figure S2). One protein’s solubility (name: agaW, N-acetylgalactosamine-specific enzyme IIC component of PTS) was improved to 0.9951 from 0.1 after adding the designed short peptide tags. When we allowed adding only 20 instead of 30 amino acids, the improvement of predicted solubility slightly decreased (Supplementary Figure S2). Since it is easier and cheaper to add 20 amino acids in experiments than 30, we adopted adding 20 amino acids as the constraint in the rest of this study.

To make this study more relevant to the imperative applications of recombinant enzymes, we selected six proteins which were important in engineering metabolic pathway of *E. coli* to produce valuable metabolites (information of these proteins is provided in caption of Figure 2). These proteins’ predicted solubility was lower than 0.6. Adding 20 amino acids also substantially improved the predicted solubility of all the six proteins (Figure 2). Three proteins (tal, dxs and valC) were chosen to experimentally validate the optimization results since their original predicted solubility was low and the predicted solubility was substantially improved through the optimization.

**Figure 2.**
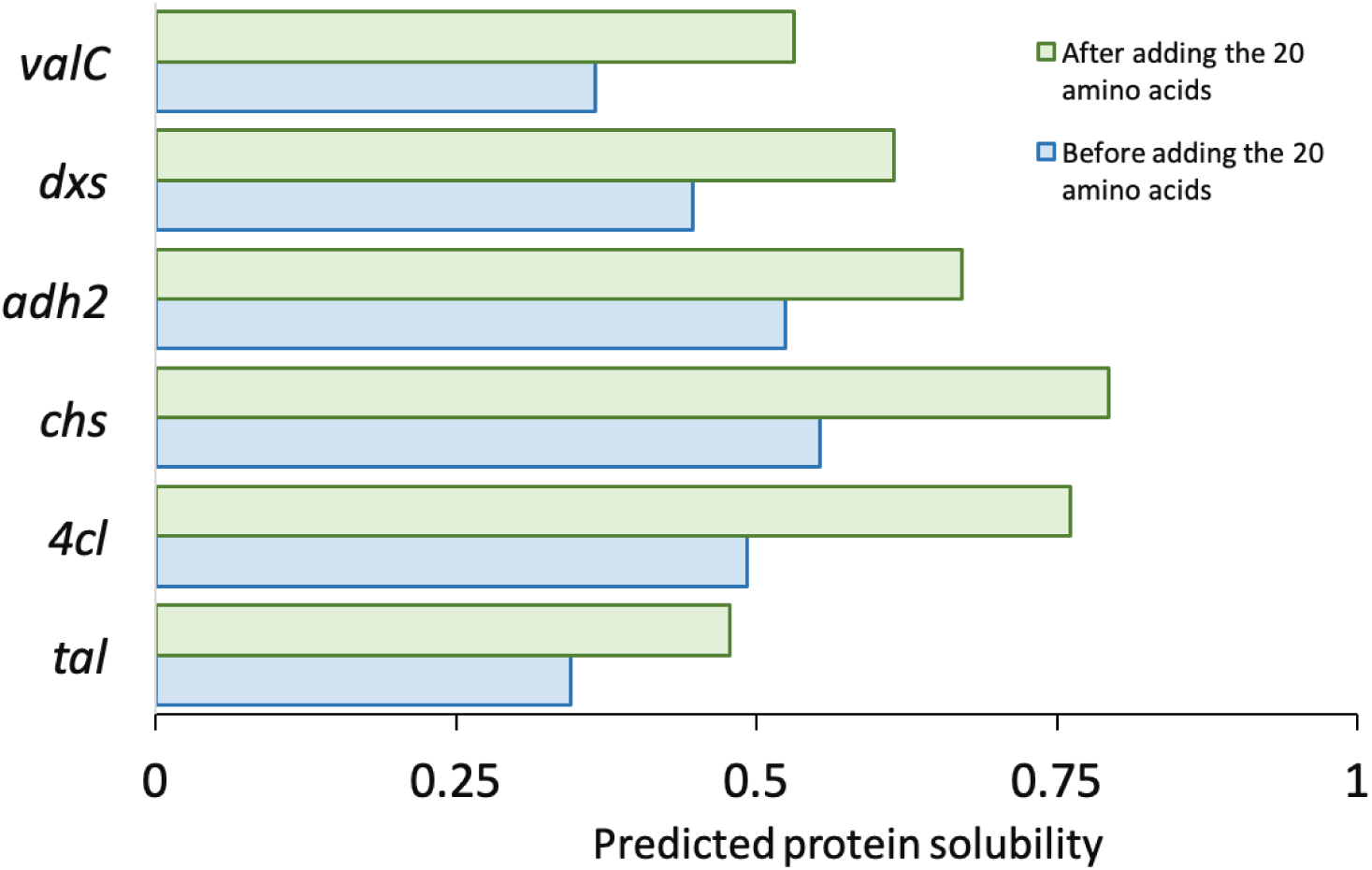
The predicted solubility before and after adding 20 amino acids for six proteins commonly used by our laboratory. The six proteins were valC (valencene synthase), dxs (1-deoxy-D-xylulose-5-phosphate synthase), adh2 (alcohol dehydrogenase), chs (chalcone synthase), 4cl (4-coumarate-CoA ligase) and tal (tyrosine ammonia-lyase). Their sequences were listed in Supplementary Table S7. Before adding the tags, the protein solubility of them was predicted by SVM and recorded. Then GA was used to optimize their solubility by adding 20 amino acids. The protein solubility after adding the tags was also recorded for comparison.

We also included agaw in the test because of the large improvement we observed in the *in silico* optimization. The number of the amino acids to be added was allowed to be decimal during the optimization and was rounded for experimental validation. The predicted solubility after rounding the number of the amino acids added was very similar to that before rounding for all the tested proteins (Supplementary Table S6). To generate sequence of the two tags to be added to a protein from the number of amino acids we minimized the occurrence of amino acid repeats, which reduced the difficulty in synthesizing the DNA. The sequence of the tags for those four proteins is listed in the Supplementary Table S7.

### Experimental validation of the optimized protein sequence

We constructed expression vectors to express the four proteins with and without the optimized tags. Among them, protein agaw cannot be expressed (as determined by using SDS-PAGE) with and without the tags, which may be caused by the unstable protein structure or unsuitable experimental conditions. Protein valC can be expressed only without the peptide tags which may have impaired the protein stability. Protein tal and dxs were expressed with and without the tags (Figure 3). Protein solubility of tal and dxs was increased by 118% and 16% respectively by adding the tags.

**Figure 3.**
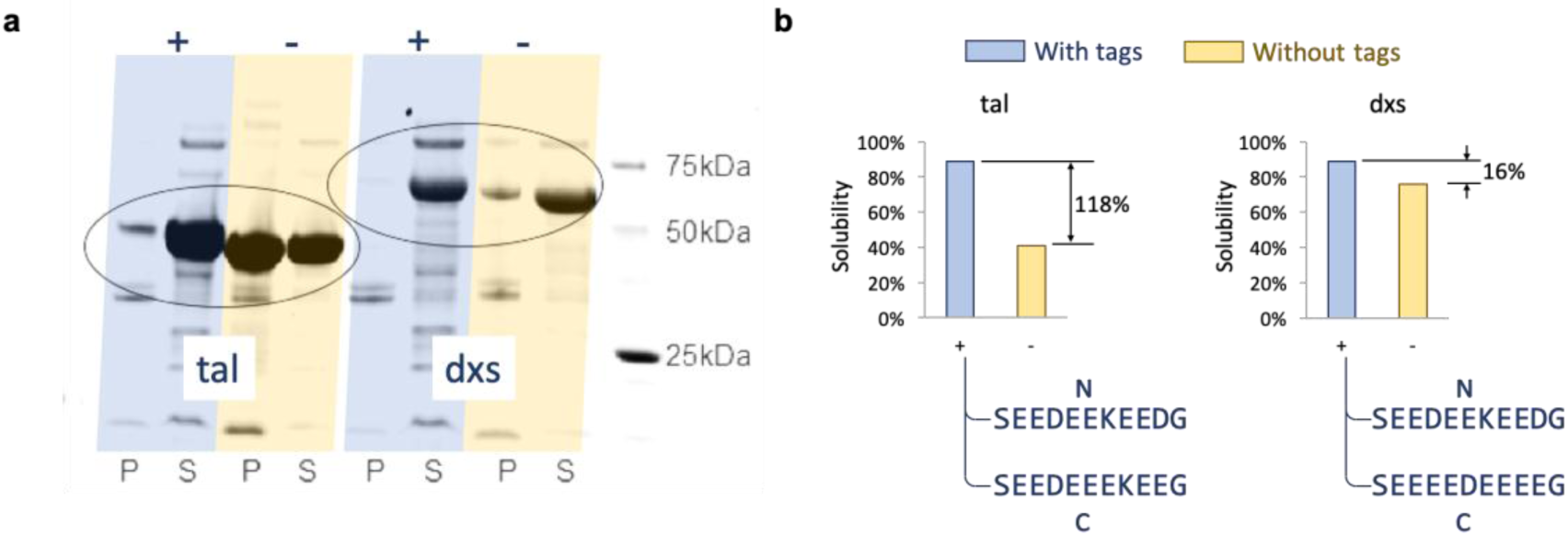
(**a)** The SDS-PAGE analysis of protein tal and dxs expressed in *E. coli* with and without tags designed by our optimization algorithm. “+” and “-” represented expressed proteins with and without peptide tags respectively. “P” and “S” represented the pellet fraction (insoluble) and supernatant fraction (soluble), respectively. The oval shapes highlight the bands of dxs and tal proteins. Protein tal and dxs were expressed in K3 medium with 20 g/L glucose at 30 °C. (**b)** Quantitative presentation of the SDS-PAGE images in **a**. The protein solubility was the ratio of soluble protein amount to the total protein amount. The protein amount was estimated by using band intensity on SDS-PAGE images. The sequences of the designed tags for N-terminal and C-terminal were shown. The amino acid S and G on the two ends of the tags were the linkers for GT DNA assembly standard, which was used to construct the plasmids in this study^40^.

By observing the amino acids added to dxs and tal (Figure 3b and Supplementary table 5), it can be found that their peptide tags were dominated by aspartic acid (D) and glutamic acid (E). Aspartic acid and glutamic acid are the two negatively charged amino acids among the 20 amino acids. Adding them may introduce repulsive electrostatic interactions between protein molecules to prevent aggregation and to provide sufficient time for correct folding of proteins^27^. The similarity of the peptide tags inspired us to test whether one tag designed for one protein can be used to improve solubility of another protein. We found that the tags optimized for improving solubility of tal could also increase both predicted and measured solubility of dxs, and vice versa (Figure 4a). Another protein (name: ada, aldehyde dehydrogenase) used in a project of our laboratory was also tested with the tag designed for tal and its predicted and measured solubility were also enhanced (Figure 4a). The results of switching tags suggested that the tags we designed may be generally effective in improving protein solubility.

**Figure 4.**
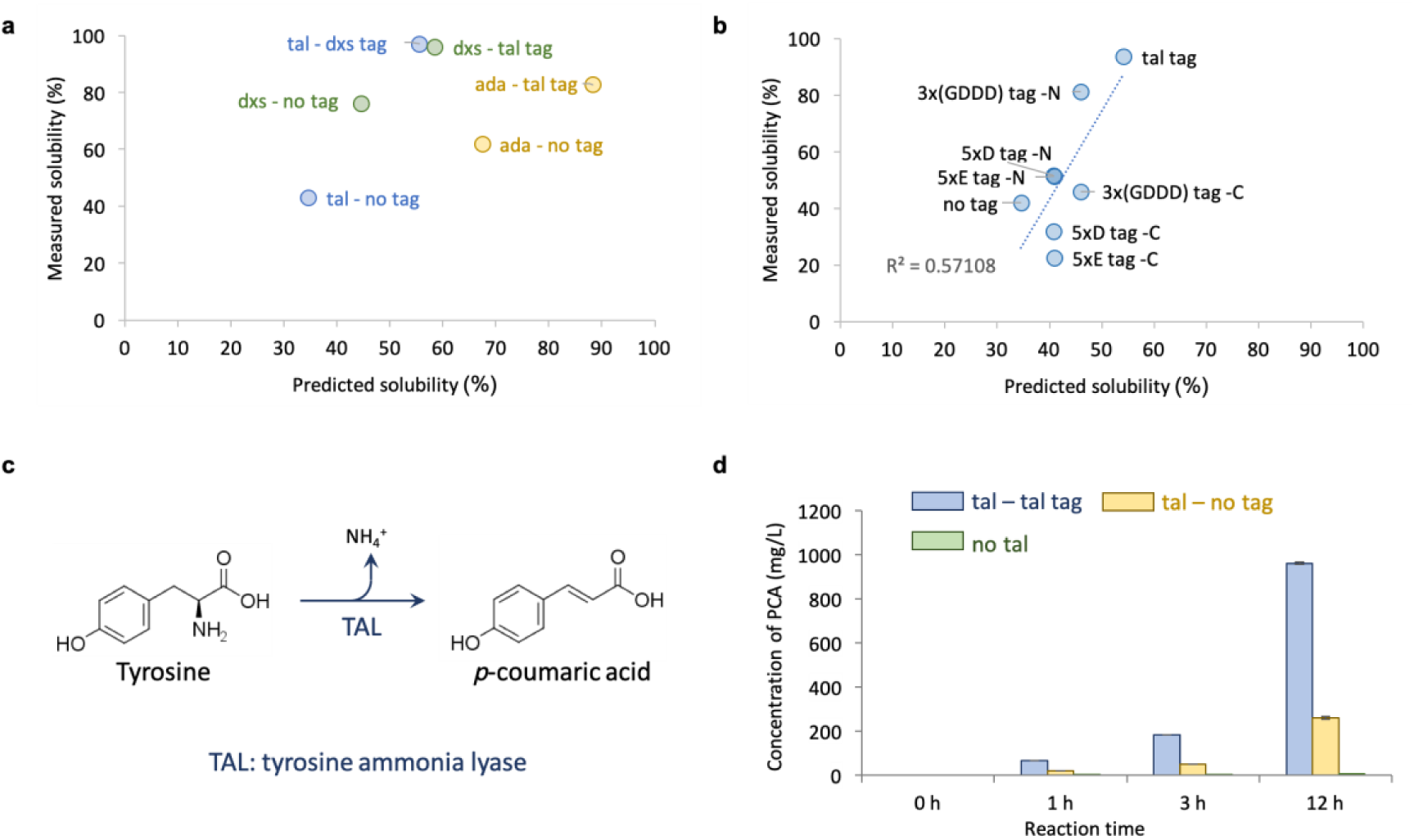
(**a**) The predicted and measured solubility of tal, dxs and ada after adding tags designed for other proteins. The purpose of switching tags for proteins was to test if the solubility-enhancing tags are generally effective in improving protein solubility. The same protein was labelled by using the same color to highlight the data before and after adding tags. In the data labels, the text before “-” indicates protein name and the text after “-” indicates the tags used if any. In the process of measuring the solubility, the protein expression condition was K3 medium with 20 g/L glucose at 30 °C. (**b**) The comparison of the tags designed in this study with tags used in previous studies. Protein tal was the only model protein used in this plot. No tag: solubility of tal without any tag. Tal tag: solubility of tal when we added the tags that were designed by our optimization algorithm for tal. 5xE tag -N/C: solubility of tal when 5xE tag (EEEEE) was added to its N- or C-terminus. 5xD tag -N/C: solubility of tal when 5xD tag (DDDDD) was added to its N- or C-terminus. 3x(GDDD) -N/C: solubility of tal when 3x(GDDD) tag (GDDDGDDDGDDD) was added to its N- or C-terminus. 5xD, 5xE and 3x(GDDD) were three tags used in a previous study and used here for comparison^27^. Since in previous study, only one tag was added to one protein, either at N- or C-terminus, we tested both cases for each tag. The two tags we designed for tal were added to both ends of tal (Figure 1 and 3b). The sequences of all the tags are provided in Supplementary Table S7. In the process of measuring the solubility, the protein expression condition was K3 medium with 20 g/L glucose at 30 °C. (**c**) The reaction catalyzed by enzyme tal. (**d**) The protein activity of protein tal before and after introducing tal tag. The product of the reaction catalyzed by enzyme tal was p-coumaric acid (PCA) and its concentration was used to indicate the activity of protein tal. Cell lysate containing tal was used in the reaction. tal – tal tag: the strain containing tal with the tags designed in this study. Tal – no tag: the strain containing tal without any tag. No tal: the strain that did not express tal. The bars indicate the mean of six replicates. The error bars indicate standard error of six replicates.

### Protein activity also improved by the optimization

The ultimate goal of this project was to improve activity of enzymes and their solubility was used as proxy because of the aforementioned reasons. Following the success of improving protein solubility, we measured activity of tal with and without the tags. Protein tal is tyrosine ammonia lyase which can deaminate tyrosine to produce coumaric acid (Figure 4c). It is very useful in producing flavonoids by using engineered microbes^28, 29^. Tal activity was increased by 269% by adding the tags we designed for it (Figure 4d, based on 12 h reaction). The extent of the increase in activity was even larger than that in solubility, suggesting that adding the tags may also increase the expression level and/or specific activity of tal. This result proved that our optimization scheme for protein solubility was also effective for improving protein activity and using protein solubility as a proxy to increase protein activity was reasonable.

### Tags designed under more constrained conditions

Among the four proteins selected for experimental validation, the protein valC (valencene synthase) cannot be synthesized only after the tags were added. This may be caused by the fact that the stability of protein valC was damaged after adding the tags. Our prediction model and optimization algorithm only took the protein solubility into account. However, other properties of the protein may be changed during the addition of highly charged tags, such as the protein stability. Therefore, we explored whether the peptide tags including mainly aspartic acid and glutamic acid can be replaced by tags that contain less charged amino acids to improve protein solubility.

The constrained condition that the number of aspartic acid and glutamic acid cannot be more than a threshold was therefore set in the optimization algorithm. The threshold was from 0 to 10 with step size of 1 for aspartic acid and glutamic acid respectively (Supplementary Table S8). When the limitation of addition number for aspartic acid and glutamic acid was reduced gradually from 10, the predicted solubility was decreasing but the change was small. With the decrease in the number of aspartic acid and glutamic acid, the number of lysine (K) increased substantially. Other amino acids only had a relatively small increase in the optimization solutions. When the constrained condition was very strict, for example, no aspartic acid and glutamic acid were allowed, the amino acids introduced were mostly alanine (A).

Another constrained condition was explored which limited the net charge of the peptide tags. In this case, the upper bound for the absolute value of the net charge of the tag was set as 5, 4, 3, 2, 1, and 0, respectively (Supplementary Table S9) and it could be observed that the number of alanine increased most substantially with the decrease of net charge, which was consistent with the results obtained under the other constraint and supported that introducing alanine may be beneficial for the dissolution of protein or the optimization failed to find a feasible solution under such stringent constraints.

This hypothesis was tested by doing experiments. The tags with net charge 1, 3, and 5 (Supplementary table S9) were used with protein valC. These new tags did not abolish protein expression, confirming the hypothesis that excessive amount of aspartic acid and/or glutamic acid may destabilize certain proteins. However, the solubility of protein valC was not improved by the tags (Supplementary Figure S3). Protein valC may have strong affinity to cellular membranes and thus cannot be solubilized by the designed tags.

### Comparison with previous studies

To improve protein solubility, some trial-and-error procedures were developed by introducing small polyionic tags^30, 31, 32^. Small peptide tags have been used as solubility-enhancing tags for a long time because they are short and less likely to interfere with protein structure^30^. One study indicated non-polar surface and positively-charged patches contributed to the separation of the soluble and insoluble proteins^31^. It was demonstrated that a concentration of positive charge may tend towards lower folded state stability through unfavourable charge interactions and result in insolubility. In addition, a negatively charged fusion tag, NT11, was also developed to enhance protein expression in *E. coli*^32^. However, these previous studies explored tags by trial and error and cannot provide a generally useful quantitative model which can forecast performance of tags with proteins which have not been tested. Among the diverse solubility-enhancing tags that have been tested, the ones that are rich in aspartic acid and glutamic acid were also studied before^27^.

To find out if the tags we obtained from our optimization were more effective than these published ones, we compared them by using our predictive model and by conducting experiments. We used tal as the model protein here, because its solubility was experimentally confirmed to be low and its measured solubility can be substantially improved by adding tags. The results were shown in Figure 4b and protein tal without tag was used as the control. All the three previously known polyionic tags increased solubility of tal when added to tal, based on experimental measurement. But none of them out-performed the tags identified in our optimization, supporting the usefulness of the tags and the optimization procedure we reported here. In addition, there was a desirable correlation between the predicted protein solubility and measured protein solubility. The linear correlation between predicted solubility and measured solubility was quantified by R^2^ with a value of 0.57. Although the previous study exploded tags including aspartic acid and glutamic acid by trial and error, our study provided better optimization performance and a generally effective quantitative model.

## Discussions

### Using machine learning for optimizing protein properties

Using machine learning to assist the selection of proteins with specific properties has been explored recently^33, 34, 35^. Heckmann et al. utilized machine learning to predict the turnover number of enzymes in *E. coli* to optimize the growth rate, proteome composition and physiology of organisms^34^. And the prediction results were further used to parameterize two mechanistic genome-scale models more accurately. The machine learning model was trained by using the information of protein structure, biochemistry properties and assay conditions^34^, whereas protein sequences were used to train our prediction model. Therefore, their model cannot be used to optimize protein sequence for improving protein activity. Wu et al. incorporated machine learning into the directed evolution workflow to help them identify proteins with high fitness value^33^. Then it was applied to engineer an enzyme for stereodivergent carbon–silicon bond formation, a new-to-nature chemical transformation. However, their training data for machine learning only included variants mutated at four amino acid residues. A protein might include multiple positions for mutagenesis and information of four positions is not representative enough to train a machine learning model to handle other positions. The selection of mutagenesis positions need to be customized by prior knowledge on the structure of proteins. Yang et al. then reviewed the machine-learning-guided directed evolution further^35^. The different representation methods of protein sequence, prediction models, optimization methods, and the training data of machine learning models were discussed for different applications. Compared with the study mentioned above^33, 35^, we do not need to train our optimization and prediction model again when we handle a new protein. In our study, we utilized the machine learning model to identify proteins with another desired property, protein solubility. Our training dataset was obtained by using various proteins of *E. coli* and the optimization methodology did not need any customization and knowledge in biochemistry for new target proteins. With only the sequence information, our optimization model can provide effective guide for improving protein solubility and activity. In addition, rather than using mutation to improve the protein properties, we added small peptide tags to improve protein solubility and activity to avoid destroying the function of the original proteins.

### The contribution of aspartic acid and glutamic acid

In this study, we designed a novel methodology to apply a predictive model of protein solubility to improve protein solubility by adding short peptide tags. Aspartic acid and glutamic acid dominated the tags that were obtained by using our optimization strategy. This finding was consistent with the conclusion of an experiment we did to determine which amino acids were the most important in determining accuracy of our solubility-predicting model. In the experiment, we removed the percentage information of two amino acids and evaluated the negative impact on the performance of the predictive model. The model’s inputs were composition of 20 amino acids, among which the percentages of 19 amino acid were independent. As a result, removing information of only one amino acid would have no impact on model performance and we had to remove the percentages of two amino acids. We evaluated all the combinations of two amino acids. After removing aspartic acid or glutamic acid, the decrease of the prediction performance represented by R^2^ was the most substantial (Figure 5), indicating they were the most important ones for the model to be accurate. The causal relationship of the observations from this experiment and the optimization experiment could be that these two negatively charged amino acids had large positive influence on protein solubility (as seen in the optimization experiment), so they were important to the accuracy of the model prediction (as observed in the importance analysis experiment). In addition, arginine which also showed some influence on the prediction performance when it was removed, did not appear in the optimization results. This might be caused by that arginine negatively affected the protein solubility and this hypothesis was tested (Supplementary Figure S4). After adding 20 arginines to the six proteins from our laboratory, all the predicted solubility was decreased. The suspected effects of glutamic acid, aspartic acid and arginine were also supported by their spearman correlation coefficients (Figure 5c), which were obtained by analyzing the large dataset we used to train our model. There were some amino acids that were identified to be important by spearman coefficient (Figure 5c) but were not found to be critical to model performance (Figure 5a), such as tryptophan and phenylalanine. It may be due to that spearman coefficient alone is not sufficient to quantitatively describe the effects of amino acid on protein solubility because of its qualitative nature and it did not consider abundance of other amino acids (Figure 5b). In this study, we have shown that our machine learning model is able to quantitatively describe the relationship and guide optimization of protein sequence.

**Figure 5.**
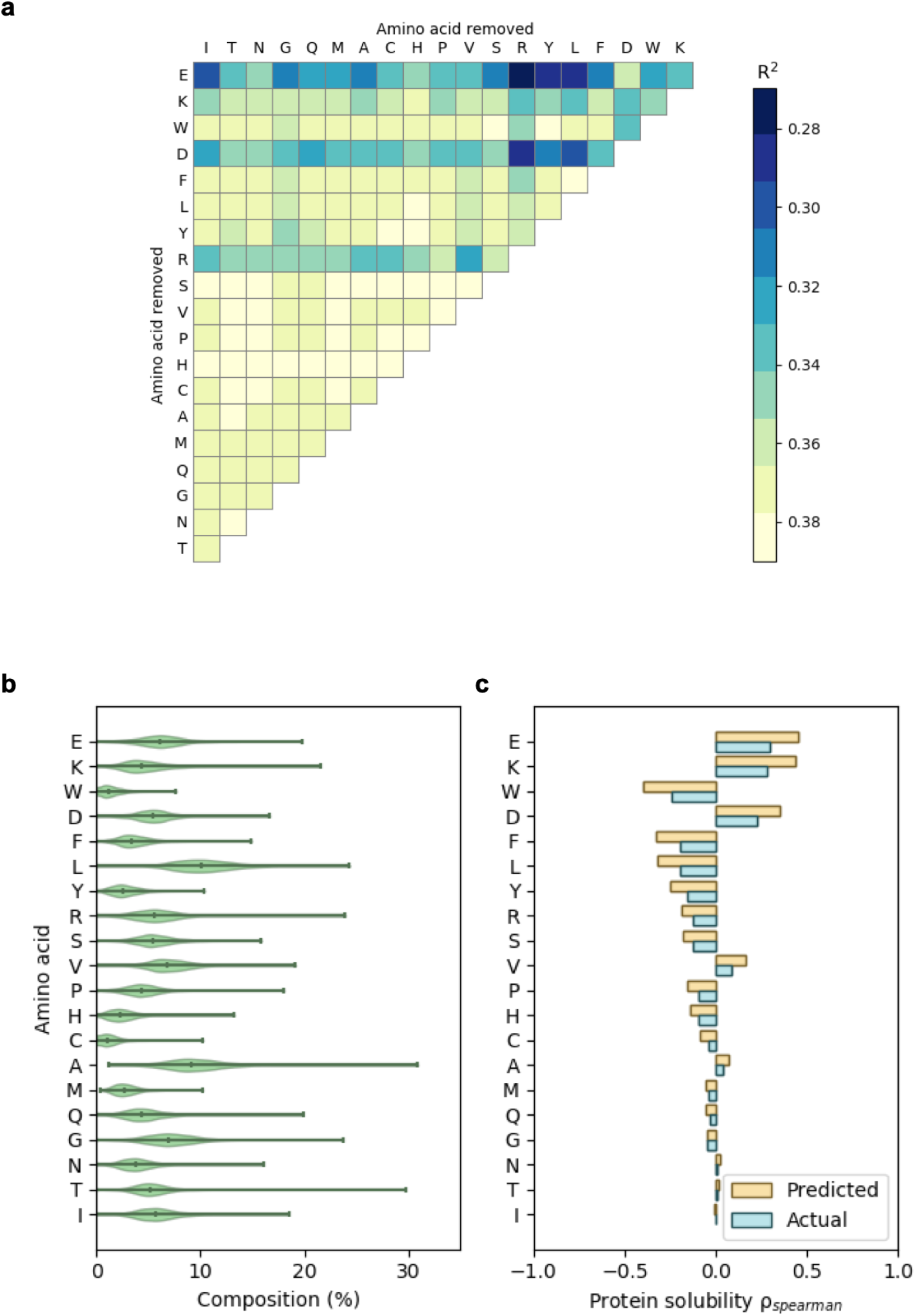
(**a**) Importance of various amino acids in determining the accuracy of the SVM regression model. The R^2^ of the SVM model was shown by using a heat map after removing the information of two types of amino acids. Model training is described in Materials and Methods. Single letter amino acid abbreviations are used in this figure. All the combinations of removing two types of amino acids are tested and the performance of the resulting models is presented in the upper triangular matrix. Performance of the models was gauged by using R^2^, which is presented here by using color (a color bar is provided). The darker the color is, the more important the related amino acids are to the model performance. (**b**) The distribution of amino acid composition (the input variables of the SVM model we used) among all the proteins in the eSol database (the date source we used to train the SVM model). The violin plot showed the mean value and the range of the amino acid composition used to train the SVM model. (**c**) The Spearman’s rank correlation between actual/predicted protein solubility and various amino acids. Spearman’s correlation, *ρ*_*spearman*_, is a measure of monotonicity and represents the general sensitivity of solubility to amino acid composition. A comparison between the Spearman’s rank correlation tornado plot for actual solubility and predicted solubility depicted how the model captured and magnified general trends between amino acid composition and solubility. For example, for both the actual and predicted solubility of proteins in the eSol dataset, the composition of D, E, or K was positively correlated with solubility.

When we trained the solubility-predicting model through machine learning, we did not use any biochemistry knowledge. The optimization of protein tag to maximize protein solubility was also purely mathematical without any dependence on prior knowledge. Yet, the identified most beneficial amino acids and their influence on protein solubility can be explained by using known biochemistry knowledge (electrostatic repulsion). As to why the best tags were dominated by negatively charged amino acids rather than positively charged ones, the reason might be that positively charged amino acids may also improve protein solubility but their influence is less than those of negatively charged amino acids. When the number of the negatively charged amino acids was constrained, the optimization algorithm used positively charged amino acid (lysine) to improve protein solubility, which led to less improvement in solubility than using the negatively charged ones (Supplementary Table 8 and 9).

## Methods

### Protein database

All the information of protein solubility used in our study is from the eSol database^11^ which is a unique database containing continuous values of protein solubility. After removing items without sequence information according to the previous study^26^, 3,148 proteins in the eSol dataset were used for this study. In the study which generated the dataset, the values of protein solubility were measured by synthesizing the recombinant proteins by cell-free protein expression technology and then being separated into soluble and insoluble fractions with centrifugation^11^. Solubility was defined as the ratio of supernatant protein to total protein which was quantified by SDS-PAGE.

### Training flowsheet

The whole process of rationally engineering proteins with higher solubility includes data pre-processing, training the SVM prediction model, constructing an optimization methodology, and validating the methodology. As the first step, amino acid composition was extracted from protein sequences by using Amino Acid Composition Descriptor in protr package^36^ within R software, which converted characters of amino acids into numerical values indicating amino acid composition. For the second part, the SVM model was built in MATLAB and trained following the same procedure described in the previous study^26^. Then SVM was trained with the whole dataset to predict continuous values of protein solubility from amino acid composition. For the third step, we filtered out a total of 58 proteins with low solubility of value 0.1 in the original dataset and 58 proteins were picked out. Proteins with long sequences are more challenging to synthesize in experiments, therefore the protein sequences were further filtered to have less than 333.3 amino acids (1kb), which excluded 27 proteins from the eSol database. Among the 27 proteins, the one with the minimum difference between the predicted value and the real value of protein solubility, named glcE, was selected as the sample protein to build a methodology for further optimizing protein solubility. Genetic algorithm (GA), an optimization method, was explored to search for maximum predicted solubility with constraints for the sample protein. The difference between protein solubility before and after mutagenesis was used to evaluate the optimization effect on protein solubility. Moreover, besides the sample protein, 10 proteins with solubility of value 0.1 which have the least differences between predicted and original solubility among the 27 proteins mentioned above were selected for applying the optimization methodology. Six proteins commonly used in our laboratory were also investigated for the optimization of protein solubility. Finally, among the 16 proteins selected for optimization, 4 proteins that bear low solubility before adding the tags and high predicted solubility after adding the tags were chosen for experimental validation. The original and mutated protein sequences were synthesized to validate the change of protein solubility by measuring the protein solubility with SDS-PAGE.

### Machine learning models

The regression version of SVM used in this study could also be named support vector regression (SVR) ^37^. The aim of SVR is to solve^38^

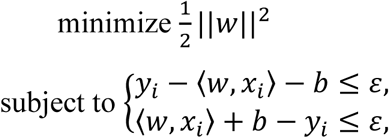

where *x*_*i*_ is a training sample with target value *y*_*i*_ and *w* is the normal vector to the hyperplane. The inner product plus intercept ⟨*w, x*_*i*_⟩ + *b* is the prediction value for that sample. The difference of predicted values and true values for targets have to be within an *ε* range, which is a parameter serving as a threshold.

A regression machine learning model SVM in MATLAB was used for optimizing protein solubility for the all the proteins in our study and was validated by experiments (Supplementary Table S10). The improved SVM model was used to optimize all the proteins *in silico* and compared with the previous one in the Discussion.

### Optimization algorithms

Genetic algorithm (GA), one of the evolutionary algorithms, is inspired by the process of natural selection observed in nature^39^. It is a frequently utilized randomized optimization algorithm for searching optima with constrained conditions. GA essentially simulates the way in which life evolves to find solutions to real world problems. In GA, the solutions to a problem are represented as a population of chromosomes evolving through successive generations. The offspring chromosomes are generated by merging two parent chromosomes by crossover or modifying a chromosome by mutation. The offspring chromosomes are evaluated according to the fitness or objective function in each generation. Chromosomes with higher fitness values have higher possibility to survive and the process will stop when the offspring chromosomes are almost identical or the terminal conditions set are reached. Strong individuals will dominate the generation through many iterations in the process with mutation, crossover and selection. The final chromosome represents an optimal or near-optimal solution for the optimization problem. In our problem, the chromosomes are the sequence of peptide tags and the fitness function is the predicted solubility for proteins after adding tags. Several hyperparameters can be tuned for the optimization algorithm, such as the population size, the number of iterations for evolution and the number of individuals mutating in each generation. We used a MATLAB Toolbox to implement the optimization (iteration number = 1,000, other parameters are provided in Supplementary Table S1). The generic structure of GA in our study can be described as follows:

~~~
**begin**:
  initiate a tag representing by a 20-dimensional vector with constrained
  conditions (sum of the vector is 20 and the value of each dimension is within range 0-20);
  evaluate the protein sequence after adding the tag;
    **while** (if termination conditions are not met):
        do crossover and mutate parent tag sequences to yield offspring sequences;
        evaluate the protein solubility for the proteins with offspring sequences;
        select and generate offspring sequence with higher solubility;
    **end while**;
**end**.
~~~

### Data visualization

The heat map was plotted by using the cmap function of the matplotlib package in Python with the values of R^2^ after removing the information of two types of amino acids. The violin plot of the amino acid compositions was made by using the violinplot function of the seaborn package in Python. Violin plot featured a kernel density estimation of the underlying distribution. Spearman’s rank correlation between amino acid composition and solubility was computed using the spearmanr function of the scipy.stats package in Python. The equation used was

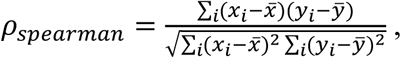

where the subscript *i* denoted the ranks, and *x* and *y* represented amino acid composition and solubility respectively.

### Chemicals in experimental validation

All chemicals were purchased from Sigma-Aldrich unless otherwise stated. All reagents used were of analytical grade. The DNA oligomers used in this study were synthesized from Integrated DNA Technologies.

### Plasmid construction

All the plasmids used in this work were constructed by using GT DNA standard^40^ (Supplementary Table S7).

### Cell culture and SDS-PAGE analysis of protein solubility

Each of constructed plasmid was introduced into *E. coli* BL21 (DE3) (C2530H, New England Biolabs) for SDS-PAGE analysis by using standard heat shock protocol. In order to test the resulting strains, single colony was inoculated into 1 mL of LB with 100 µg/mL of ampicillin, and was cultured overnight at 37 °C/250 rpm. Fifty microliters of the overnight grown cell suspension were inoculated into 5 mL of K3 medium^40^ with 100 µg/mL of ampicillin. When cell was grown to 0.4-0.6 optical density (OD) at 600, isopropyl β-D-1-thiogalactopyranoside (IPTG) was added to a final concentration of 0.1 mM. After incubated overnight at 30 °C/250 rpm, the cell culture broth was diluted to OD600 = 2.0, and centrifuged at 5000 g, 10 min. The obtained cell pellets were resuspended in 100 µL B-PER II reagent (78248, Thermo Fisher Scientific). The mixtures were incubated for 15 min at room temperature with gentle rocking, and centrifuged at 16000 g for 20 min. The obtained supernatant contained soluble cell lysates. The insoluble cell pellets were resuspended in 100 µL of 2 % w/v SDS. Both soluble and insoluble cell pellets were analyzed by using SDS-PAGE (Mini-PROTEAN® TGX™ Precast Protein Gels, 4561083, Bio-Rad). The image of the gel was processed and quantified by Gel Doc EZ Gel Documentation System (Bio-Rad).

### Tal activity assay *in vitro*

One milliliter of obtained supernatant containing soluble cell lysates was added to 4 mL of PBS buffer (pH=9.0) with 1 g/L tyrosine (final concentration) in 50 mL falcon tube and incubated at 30 °C/250 rpm. Three hundred microliters of samples were taken at 0 h, 1 h, 3 h and 12 h after incubation, and mixed with 700 µL of acetonitrile to dissolve the produced *p*-courmaric acid (PCA). The mixture was incubated at 30 °C/250 rpm for 1 h, and then centrifuged at 13,500 g for 5 min. Two microliters of the obtained supernatant was analyzed by using HPLC (Agilent 1260 Infinity HPLC) based on a previously reported method^40^.

## Supplementary information

Supplementary data are available online.

## Codes availability

We present the optimization workflow as a series of notebooks hosted on GitHub (https://github.com/xiaomizhou616/optimization_protein-solubility). The workflow can be used as a template for analysis of other expression and solubility datasets.

## Acknowledgments

We acknowledge the MOE Research Scholarship, MOE Tier-1 grant (R-279-000-452-133) and NRF CRP grant (R-279-000-512-281) in Singapore. We thank Cortes-Pena Yoel Rene for providing data visualization for data distribution and Spearman’s rank correlation tornado plot.

## Author contributions

X.H. developed the optimization algorithms and statistical analyses. W.N. performed the experimental preparation and validation, and X.M designed and guided the experiments. All of them were supervised by X.W. and K.Z.. X.H. and W.N. wrote the manuscript with inputs from all the co-authors. All authors discussed the results and commented on the manuscript.

