## Supplementary Information for "Improve Protein Solubility and Activity based on Machine Learning Models"

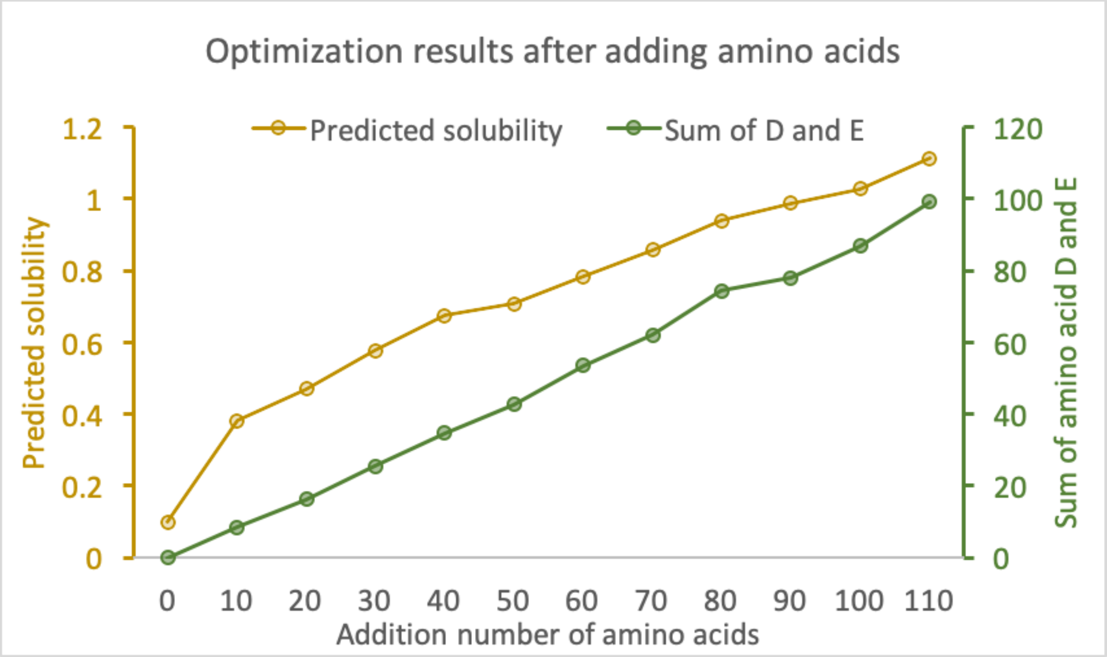


**Figure S1**. The optimization results for different lengths of peptide tags. The protein solubility of the sample protein was optimized by adding different lengths of peptide tags. The number of amino acids added on the original sequence with length 350 was searched from 0 to 110 with step size 10. The yellow and green line represented the predicted protein solubility and sum of number for amino acid D and E added respectively. Both the predicted protein solubility and sum of number for amino acid D and E added increased with the increasing addition number of amino acids. However, in reality, the extreme condition, proteins almost only containing D and E, would not exist and we should choose a reasonable number of amino acid added according to the specific situations.

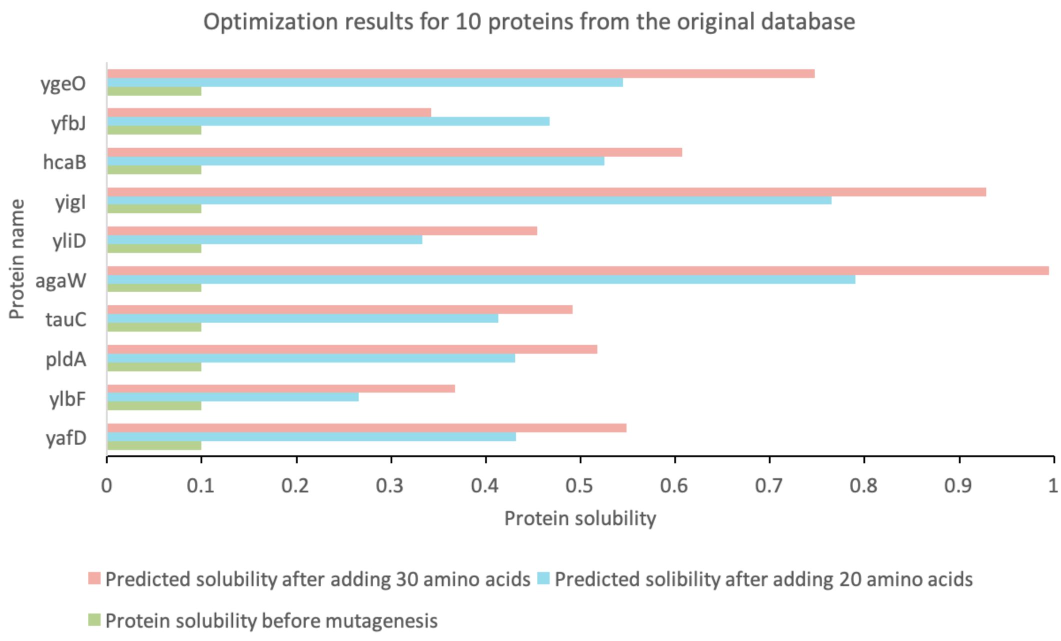


**Figure S2.** The optimization results for 10 proteins selected from the original database. The 10 proteins with solubility 0.1 from the eSol database (Niwa, et al., 2009) were optimized by adding 30 and 20 amino acids respectively. The corresponding addition number of each amino acid for the 10 proteins was recorded in the Supplementary Table S3, S4.

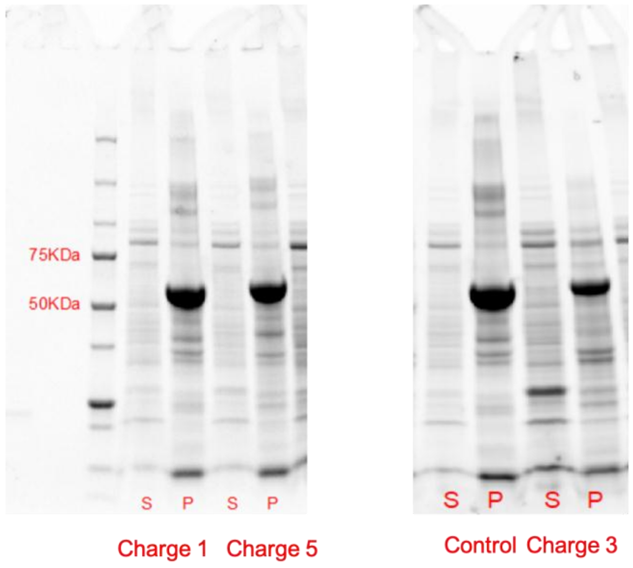


**Figure S3.** The SDS-PAGE analysis of protein valC expressed in *E.coli* with different charged tags. Protein valC was cultured in K3 medium with 20 g/L glucose at 30 ℃. “S” and “P” represented the pellet (insoluble) fraction and soluble fraction respectively. The charge labeled below indicated the tags designed in Table S9 with different charges and control is the expressed protein valC without tags.

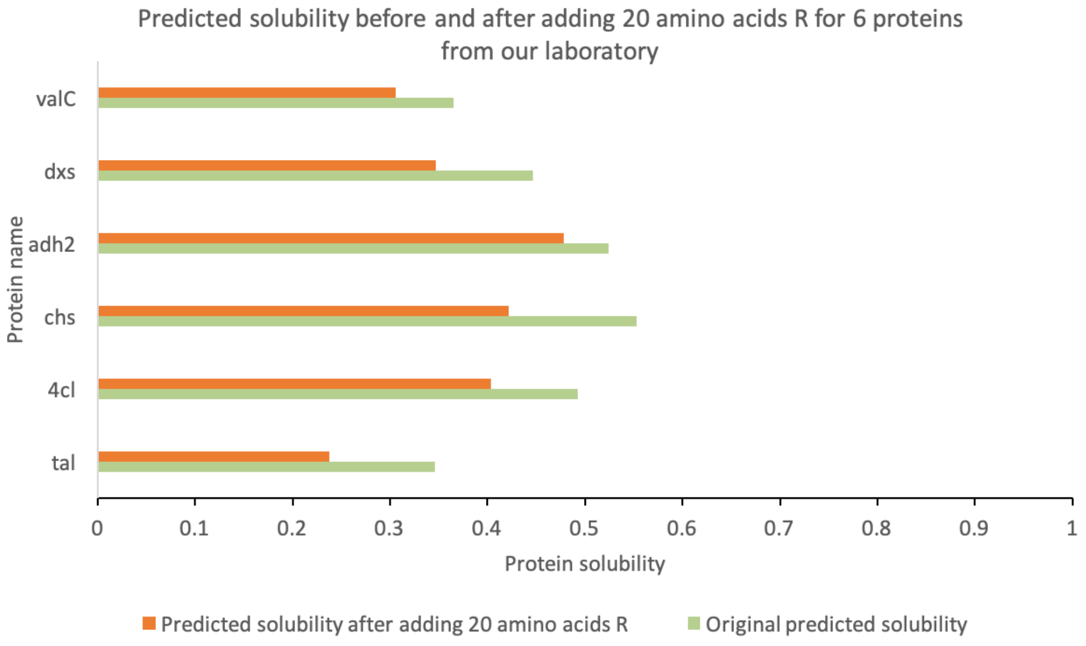


**Figure S4.** The predicted solubility before and after adding 20 amino acids for 6 proteins from our laboratory. Twenty amino acids R were added on the sequence of 6 proteins and their predicted solubility was generated from SVM model in MATLAB.

**Table S1.** Optimization results for adding 30 amino acids on sample protein

| Number | Algorithm | Amino acids added | Iteration | Population size | Variable | Predicted solubility |
|---|---|---|---|---|---|---|
| 1 | GA | 30 | 1000 | 200 | Continuous | 0.5775 |
| 2 | GA | 30 | 2000 | 200 | Integer [a] | 0.2446 |
| 3 | GA | 30 | 10000 [b] | 1000 | Integer | 0.3162 |
| 4 | GA | 30 | 10000 | 1000 | Continuous | 0.5918 |

[a] We tried to set the variables of GA as integers in MATLAB as the meaning of the variable was number of amino acid added. However, GA in MATLAB failed to deal with integer variables and equality constraints simultaneously. Due to the low efficiency and accuracy with integers here, continuous values of variables were used in the optimization algorithm and were rounded.

[b] After improving the number of iterations and population size, computing time increased substantially, while the predicted protein solubility only enhanced from 0.5775 to 0.5918. Thus, GA with default parameters has satisfied our needs and was utilized in our main work.

**Table S2**. The description of all proteins used in our study

| Name | Function |
|---|---|
| yafD | endo/exonuclease/phosphatase family protein |
| ylbF | putative anaerobic allantoin catabolic oxamate carbamoyltransferase; DUF2877 family protein |

| | |
|---|---|
| pldA | outer membrane phospholipase A |
| tauC | taurine ABC transporter permease |
| agaW | pseudo |
| yliD | putative peptide ABC transporter permease |
| yigI | 4HBT thioesterase family protein |
| hcaB | 2,3-dihydroxy-2,3-dihydrophenylpropionate dehydrogenase |
| yfbJ | 4-amino-4-deoxy-L-arabinose-phosphoundecaprenol flippase subunit ArnF |
| ygeO | pseudo |
| tal | tyrosine ammonia-lyase |
| 4cl | 4-coumarate-CoA ligase |
| chs | chalcone synthase |
| adh2 | alcohol dehydrogenase |
| dxs | 1-deoxy-D-xylulose-5-phosphate synthase |
| valC | valencene synthase |
| ada | aldehyde dehydrogenase |

**Table S3**. Addition number of amino acids for 10 samples after rounding by adding 30 amino acids

| Protein | yafD | ylbF | pldA | tauC | agaW | yliD | yigI | hcaB | yfbJ | ygeO |
|---|---|---|---|---|---|---|---|---|---|---|
| A | 0 | 0 | 0 | 0 | 0 | 0 | 0 | 0 | 0 | 0 |
| R | 0 | 0 | 0 | 0 | 0 | 0 | 0 | 0 | 0 | 0 |
| N | 0 | 0 | 0 | 0 | 0 | 0 | 0 | 0 | 0 | 0 |
| **D** | **18** | **22** | **20** | **18** | **4** | **10** | **21** | **20** | **22** | **19** |
| C | 0 | 0 | 0 | 0 | 0 | 0 | 0 | 0 | 0 | 0 |
| **E** | **8** | **4** | **7** | **7** | **24** | **17** | **7** | **6** | **2** | **8** |
| Q | 0 | 0 | 0 | 0 | 0 | 0 | 0 | 0 | 0 | 0 |
| G | 0 | 0 | 0 | 0 | 0 | 0 | 0 | 0 | 0 | 0 |
| H | 0 | 0 | 0 | 0 | 0 | 0 | 0 | 0 | 0 | 0 |
| I | 0 | 0 | 0 | 0 | 0 | 0 | 0 | 0 | 0 | 0 |
| L | 0 | 0 | 0 | 0 | 0 | 0 | 0 | 0 | 0 | 0 |

| | | | | | | | | | | |
|---|---|---|---|---|---|---|---|---|---|---|
| K | 2 | 2 | 2 | 3 | 1 | 1 | 1 | 2 | 3 | 1 |
| M | 0 | 0 | 0 | 0 | 0 | 0 | 0 | 0 | 0 | 0 |
| F | 0 | 0 | 0 | 0 | 0 | 0 | 0 | 0 | 0 | 0 |
| P | 0 | 0 | 0 | 0 | 0 | 0 | 0 | 0 | 0 | 0 |
| S | 0 | 0 | 0 | 0 | 0 | 0 | 0 | 0 | 0 | 0 |
| T | 0 | 1 | 0 | 0 | 0 | 0 | 0 | 0 | 0 | 0 |
| W | 0 | 0 | 0 | 0 | 0 | 0 | 0 | 0 | 0 | 0 |
| Y | 0 | 0 | 0 | 0 | 0 | 0 | 0 | 0 | 0 | 0 |
| V | 1 | 0 | 0 | 1 | 0 | 0 | 0 | 1 | 1 | 1 |

**Table S4**. Addition number of amino acids for 10 samples after rounding by adding 20 amino acids

| Protein | yafD | ylbF | pldA | tauC | agaW | yliD | yigI | hcaB | yfbJ | ygeO |
|---|---|---|---|---|---|---|---|---|---|---|
| A | 0 | 0 | 0 | 0 | 0 | 0 | 0 | 0 | 0 | 0 |
| R | 0 | 0 | 0 | 0 | 0 | 0 | 0 | 0 | 0 | 0 |
| N | 0 | 0 | 0 | 0 | 0 | 0 | 0 | 0 | 0 | 0 |
| **D** | **14** | **5** | **3** | **3** | **1** | **13** | **9** | **1** | **13** | **15** |
| C | 0 | 0 | 0 | 1 | 0 | 0 | 0 | 0 | 0 | 0 |
| **E** | **4** | **11** | **14** | **15** | **18** | **6** | **7** | **16** | **3** | **3** |
| Q | 0 | 0 | 0 | 0 | 0 | 0 | 0 | 0 | 0 | 0 |
| G | 0 | 0 | 0 | 0 | 0 | 0 | 0 | 0 | 0 | 0 |
| H | 0 | 0 | 0 | 0 | 0 | 0 | 0 | 0 | 0 | 0 |
| I | 0 | 0 | 0 | 0 | 0 | 0 | 0 | 0 | 0 | 0 |
| L | 0 | 0 | 0 | 0 | 0 | 0 | 0 | 0 | 0 | 0 |
| K | 1 | 2 | 1 | 0 | 0 | 0 | 3 | 1 | 2 | 1 |
| M | 0 | 0 | 0 | 0 | 0 | 0 | 0 | 0 | 0 | 0 |
| F | 0 | 0 | 0 | 0 | 0 | 0 | 0 | 0 | 0 | 0 |
| P | 0 | 0 | 0 | 0 | 0 | 0 | 0 | 0 | 0 | 0 |
| S | 0 | 0 | 0 | 0 | 0 | 0 | 0 | 0 | 0 | 0 |
| T | 0 | 0 | 0 | 0 | 0 | 0 | 0 | 0 | 0 | 0 |
| W | 0 | 0 | 0 | 0 | 0 | 0 | 0 | 0 | 0 | 0 |
| Y | 0 | 0 | 0 | 0 | 0 | 0 | 0 | 0 | 0 | 0 |
| V | 0 | 1 | 0 | 0 | 0 | 0 | 0 | 0 | 1 | 1 |

**Table S5**. Addition number of amino acids for 6 proteins in our laboratory after rounding by adding 20 amino acids

| Protein | tal | 4cl | chs | adh2 | dxs | valC |
|---|---|---|---|---|---|---|
| A | 0 | 0 | 0 | 0 | 0 | 0 |

| | | | | | | |
|---|---|---|---|---|---|---|
| R | 0 | 0 | 0 | 0 | 0 | 0 |
| N | 0 | 0 | 0 | 0 | 0 | 0 |
| **D** | **4** | **3** | **4** | **13** | **2** | **14** |
| C | 0 | 0 | 0 | 0 | 0 | 0 |
| **E** | **12** | **15** | **14** | **5** | **15** | **3** |
| Q | 0 | 0 | 0 | 0 | 0 | 0 |
| G | 0 | 0 | 0 | 0 | 0 | 0 |
| H | 0 | 0 | 0 | 0 | 0 | 0 |
| I | 0 | 0 | 0 | 0 | 0 | 0 |
| L | 0 | 0 | 0 | 0 | 0 | 0 |
| K | 2 | 1 | 0 | 1 | 1 | 0 |
| M | 0 | 0 | 0 | 0 | 0 | 0 |
| F | 0 | 0 | 0 | 0 | 0 | 0 |
| P | 0 | 0 | 0 | 0 | 0 | 0 |
| S | 0 | 0 | 0 | 0 | 0 | 0 |
| T | 0 | 0 | 0 | 0 | 0 | 1 |
| W | 0 | 0 | 0 | 0 | 0 | 0 |
| Y | 0 | 0 | 0 | 0 | 0 | 0 |
| V | 0 | 1 | 1 | 0 | 0 | 0 |

**Table S6**. The predicted protein solubility before and after rounding the number of amino acids added

| Protein | Solubility after rounding | Solubility before rounding |
|---|---|---|
| yafD | 0.4383 | 0.4319 |
| ylbF | 0.2675 | 0.266 |
| pldA | 0.4252 | 0.4305 |
| tauC | 0.4071 | 0.4131 |
| agaW | 0.804 | 0.7901 |
| yliD | 0.3345 | 0.3331 |
| yigI | 0.7592 | 0.7647 |
| hcaB | 0.5201 | 0.5256 |
| yfbJ | 0.4651 | 0.4676 |
| ygeO | 0.5698 | 0.5446 |
| tal | 0.4737 | 0.4777 |
| 4cl | 0.7647 | 0.7606 |
| chs | 0.7876 | 0.7925 |
| adh2 | 0.6704 | 0.6703 |
| dxs | 0.6081 | 0.6141 |

| | | |
|---|---|---|
| valC | 0.5708 | 0.531 |

**Table S7**. The sequence of peptide tags tested by experiments

See attached excel "Supplementary Table 7.xlsx".

**Table S8**. Addition number of amino acids for ValC after rounding by limiting D and E added

| Limitation for D and E | 10 | 9 | 8 | 7 | 6 | 5 | 4 | 3 | 2 | 1 | 0 |
|---|---|---|---|---|---|---|---|---|---|---|---|
| A | 0 | 0 | 0 | 0 | 0 | 0 | 0 | 0 | 0 | 1 | 20 |
| R | 0 | 0 | 0 | 0 | 0 | 0 | 0 | 0 | 0 | 0 | 0 |
| N | 0 | 0 | 1 | 0 | 0 | 0 | 0 | 0 | 0 | 0 | 0 |
| D | 5 | 5 | 5 | 3 | 3 | 5 | 4 | 3 | 2 | 1 | 0 |
| C | 0 | 0 | 0 | 1 | 0 | 0 | 0 | 0 | 0 | 0 | 0 |
| E | 10 | 9 | 8 | 7 | 5 | 3 | 4 | 3 | 2 | 1 | 0 |
| Q | 0 | 0 | 0 | 0 | 0 | 0 | 0 | 0 | 0 | 0 | 0 |
| G | 0 | 0 | 0 | 0 | 0 | 0 | 0 | 0 | 0 | 0 | 0 |
| H | 0 | 0 | 0 | 0 | 0 | 0 | 0 | 0 | 0 | 0 | 0 |
| I | 0 | 0 | 0 | 2 | 0 | 0 | 0 | 0 | 0 | 0 | 0 |
| L | 0 | 0 | 0 | 0 | 0 | 0 | 0 | 0 | 0 | 0 | 0 |
| **K** | **2** | **3** | **2** | **2** | **11** | **10** | **9** | **11** | **12** | **11** | **0** |
| M | 0 | 0 | 1 | 2 | 0 | 0 | 0 | 1 | 1 | 3 | 0 |
| F | 0 | 0 | 0 | 0 | 0 | 0 | 0 | 0 | 0 | 0 | 0 |
| P | 0 | 0 | 0 | 0 | 0 | 0 | 0 | 0 | 0 | 0 | 0 |
| S | 0 | 0 | 0 | 0 | 0 | 0 | 0 | 0 | 0 | 0 | 0 |
| T | 0 | 0 | 1 | 0 | 0 | 0 | 1 | 0 | 0 | 0 | 0 |
| W | 0 | 0 | 0 | 0 | 0 | 0 | 0 | 0 | 0 | 0 | 0 |
| Y | 0 | 0 | 0 | 0 | 0 | 0 | 0 | 0 | 0 | 0 | 0 |
| V | 2 | 1 | 1 | 1 | 0 | 1 | 1 | 1 | 1 | 1 | 0 |
| solubility | 0.5376 | 0.5344 | 0.5250 | 0.5095 | 0.5259 | 0.5248 | 0.5215 | 0.5203 | 0.5148 | 0.5017 | 0.4449 |

**Table S9**. Addition number of amino acids after rounding for ValC by limiting net charge added

| Limitation for net charge | 5 | 4 | 3 | 2 | 1 | 0 |
|---|---|---|---|---|---|---|
| **A** | **1** | **3** | **6** | **12** | **16** | **20** |
| R | 0 | 0 | 0 | 0 | 0 | 0 |
| N | 0 | 0 | 0 | 0 | 0 | 0 |
| **D** | **5** | **4** | **1** | **2** | **1** | **0** |

| | | | | | | |
|---|---|---|---|---|---|---|
| C | 3 | 4 | 0 | 0 | 0 | 0 |
| E | 0 | 0 | 1 | 0 | 0 | 0 |
| Q | 0 | 1 | 0 | 0 | 0 | 0 |
| G | 0 | 0 | 0 | 0 | 0 | 0 |
| H | 0 | 0 | 0 | 0 | 0 | 0 |
| I | 1 | 0 | 0 | 1 | 0 | 0 |
| L | 0 | 0 | 0 | 0 | 0 | 0 |
| K | 0 | 0 | 1 | 0 | 0 | 0 |
| M | 1 | 1 | 0 | 0 | 0 | 0 |
| F | 0 | 0 | 0 | 0 | 0 | 0 |
| P | 0 | 0 | 0 | 0 | 1 | 0 |
| S | 0 | 0 | 0 | 0 | 0 | 0 |
| T | 5 | 4 | 0 | 0 | 1 | 0 |
| W | 0 | 0 | 0 | 0 | 0 | 0 |
| Y | 0 | 0 | 0 | 0 | 0 | 0 |
| V | 2 | 4 | 10 | 3 | 1 | 0 |
| solubility | 0.5135 | 0.5093 | 0.5063 | 0.4878 | 0.4805 | 0.4725 |

**Table S10**. Optimization for prediction model SVM [a]

| Model | Kernel | BoxCon-straint | KernelScale | Epsilon | $R^2$ | MSE | Optimization method |
|---|---|---|---|---|---|---|---|
| SVM1 | linear | 0.0115 | 0.0022 | 0.0022 | 0.2202 | 0.0802 | default tool |
| SVM2 | rbf | 742.2979 | 0.9629 | 0.2308 | 0.3832 | 0.0635 | GA |

[a] To further improve our methodology, we consider enhancing the prediction model SVM whose outputs are the value of the objective function of GA. The hyperparameter tuning method of the SVM model in MATLAB was not very effective and we tried to use GA to optimize the three hyperparameters with the customized objective function R2 (Supplementary Table S10). After optimization, the prediction was improved according to the value of R2 (from 0.2202 to 0.3832) and MSE (from 0.0802 to 0.0635). Then the optimization approach with the improved SVM was applied to the 16 proteins by adding 20 amino acids. The optimization results reflected the same trend with the optimization approach using the previous SVM model. As shown in Supplementary Table S11 and S13, the amino acids added are still mainly D and E. In addition, the protein with the highest predicted solubility in the original database, agaw, is still the most promising one among the 10 proteins according to Supplementary Table S11. According to the definition of the protein solubility in our study, the value of protein solubility should not be higher than 1 but the model cannot interpret the biological meaning. It might be possible that predicted solubility is more than 1 when the mutated protein

sequence is rich in D and E. Moreover, the protein with lowest predicted solubility after optimization is still ylbF. In the Supplementary Table S12, the proteins with highest predicted solubility and lowest predicted solubility were still chs and valC respectively. On the other hand, there are also differences between the optimization results in Supplementary Table S4, S5 and S11, S12. For the same protein, the predicted solubility and the number of amino acids added changed slightly. For example, protein tal and 4cl need more amino acid D than E in Supplementary Table S12, whereas the solutions include more amino acid E than D in Supplementary Table S5. This might be caused by two reasons. At first, since the GA is a random-based evolutionary algorithm where the solutions might be slightly different even with the same hyperparameters. Both adding amino acid D or E will increase the predicted solubility and the solutions of GA sometimes reach more D and sometimes more E. The second guess is that there is some complex mechanism behind and the need for amino acid D and E for different proteins might be different. The performance of prediction model does not influence the optimization solutions substantially for our problem and the experimental validation with the previous SVM model is still meaningful. For the further work, more experimental validation could be done to test whether this improved SVM model can provide more accurate guide in applications.

**Table S11**. Optimization for 10 samples by adding 20 amino acids after improving prediction model

| Protein | yafD | ylbF | pldA | tauC | agaW | yliD | yigI | hcaB | yfbJ | ygeO |
|---|---|---|---|---|---|---|---|---|---|---|
| A | 0 | 0 | 0 | 0 | 0 | 1 | 0 | 0 | 0 | 0 |
| R | 0 | 0 | 0 | 0 | 0 | 0 | 0 | 0 | 0 | 0 |
| N | 0 | 0 | 0 | 0 | 0 | 0 | 0 | 1 | 0 | 0 |
| **D** | 15 | 17 | 0 | 0 | 2 | 3 | 0 | 1 | 0 | 0 |
| C | 0 | 0 | 0 | 0 | 0 | 0 | 0 | 0 | 0 | 0 |
| **E** | 3 | 2 | 17 | 18 | 17 | 13 | 18 | 17 | 16 | 15 |
| Q | 0 | 0 | 0 | 0 | 0 | 0 | 0 | 0 | 0 | 0 |
| G | 0 | 0 | 0 | 0 | 0 | 0 | 0 | 0 | 0 | 0 |
| H | 0 | 0 | 0 | 0 | 0 | 0 | 0 | 0 | 0 | 0 |
| I | 0 | 0 | 0 | 0 | 0 | 1 | 0 | 0 | 0 | 0 |
| L | 0 | 0 | 0 | 0 | 0 | 0 | 0 | 0 | 1 | 1 |
| K | 0 | 0 | 0 | 0 | 0 | 0 | 0 | 0 | 1 | 0 |
| M | 0 | 0 | 0 | 0 | 0 | 0 | 0 | 0 | 0 | 0 |
| F | 0 | 0 | 0 | 0 | 0 | 0 | 0 | 0 | 0 | 0 |
| P | 0 | 0 | 1 | 0 | 1 | 0 | 0 | 1 | 0 | 3 |
| S | 0 | 0 | 0 | 0 | 0 | 0 | 0 | 0 | 0 | 0 |
| T | 0 | 0 | 0 | 0 | 0 | 0 | 0 | 0 | 0 | 0 |
| W | 0 | 0 | 0 | 0 | 0 | 0 | 0 | 0 | 0 | 0 |
| Y | 0 | 0 | 0 | 0 | 0 | 0 | 0 | 0 | 1 | 0 |
| V | 0 | 0 | 1 | 0 | 0 | 1 | 0 | 0 | 0 | 0 |
| Solubility | 0.5343 | 0.3422 | 0.5555 | 0.5288 | 1.0744 | 0.3859 | 0.9250 | 0.6553 | 1.0654 | 0.7413 |

**Table S12**. Optimization for 6 samples by adding 20 amino acids after improving prediction model

| Protein | tal | 4cl | chs | adh2 | dxs | valC |
|---|---|---|---|---|---|---|
| A | 0 | 0 | 0 | 0 | 0 | 0 |
| R | 0 | 0 | 0 | 0 | 0 | 0 |
| N | 0 | 0 | 0 | 0 | 0 | 0 |
| **D** | **18** | **16** | **1** | **2** | **3** | **3** |
| C | 0 | 0 | 0 | 0 | 0 | 0 |
| **E** | **1** | **1** | **16** | **14** | **13** | **15** |
| Q | 0 | 0 | 0 | 0 | 0 | 0 |
| G | 0 | 0 | 1 | 0 | 0 | 0 |
| H | 0 | 0 | 0 | 0 | 0 | 0 |
| I | 0 | 0 | 0 | 0 | 0 | 0 |
| L | 0 | 0 | 0 | 0 | 0 | 0 |
| K | 0 | 0 | 0 | 0 | 0 | 0 |
| M | 0 | 0 | 0 | 0 | 0 | 0 |
| F | 0 | 0 | 0 | 0 | 0 | 0 |
| P | 0 | 0 | 1 | 1 | 0 | 1 |
| S | 0 | 0 | 0 | 0 | 0 | 0 |
| T | 0 | 1 | 1 | 0 | 0 | 0 |
| W | 0 | 0 | 0 | 0 | 0 | 0 |
| Y | 0 | 0 | 0 | 0 | 0 | 0 |
| V | 0 | 0 | 0 | 0 | 1 | 0 |
| Solubility | 0.6024 | 0.6445 | 0.7902 | 0.7332 | 0.5878 | 0.4848 |
